# Cdc42 couples septin recruitment to the axial landmark assembly via Axl2 in budding yeast

**DOI:** 10.1101/2023.08.25.554823

**Authors:** Pil Jung Kang, Rachel Mullner, Kendra Lian, Hay-Oak Park

## Abstract

Cell polarization generally occurs along a single axis that is directed by a spatial cue. Cells of the budding yeast *Saccharomyces cerevisiae* undergo polarized growth and oriented cell division in a spatial pattern by selecting a specific bud site. Haploid **a** or α cells bud in the axial pattern in response to a transient landmark that includes Bud3, Bud4, Axl1, and Axl2. Septins, a family of filament-forming GTP-binding proteins, are also involved in axial budding and recruited to an incipient bud site, but the mechanism of recruitment remains unclear. Here, we show that Axl2 interacts with Bud3 and the Cdc42 GTPase in its GTP-bound state. Axl2 also interacts with Cdc10, a septin subunit, promoting efficient recruitment of septins near the cell division site. Furthermore, a *cdc42* mutant defective in the axial budding pattern at a semi-permissive temperature had a reduced interaction with Axl2 and compromised septin recruitment in the G1 phase. We thus propose that active Cdc42 brings Axl2 to the Bud3-Bud4 complex and that Axl2 then interacts with Cdc10, linking septin recruitment to the axial landmark.

## INTRODUCTION

Cdc42 plays a central role in polarity establishment in yeast and animals. In budding yeast, Cdc42 is involved in the selection of a bud site and polarized organization of the actin and septin cytoskeletons (Bi and Park, 2012). Septins are recruited to an incipient bud site and then converted to a ring at the mother-bud neck after bud emergence (Iwase et al., 2006; Okada et al., 2013), and the septin organization depends on Cdc42 polarization (Caviston et al., 2003; Gladfelter et al., 2002; Iwase et al., 2006). Earlier studies suggested a linear morphogenetic hierarchy from spatial landmarks to the cytoskeletons via the Rsr1 and Cdc42 GTPase modules (Chant and Pringle, 1995; Kang et al., 2001; Park et al., 1997). However, temporal interactions among the landmark components, Cdc42, and septins suggest more complex crosstalk between the proteins involved in polarity establishment. The septin ring recruits the axial landmark components Bud3 and Bud4 (Chant et al., 1995; Sanders and Herskowitz, 1996), and Bud3 and Bud4, in turn, are necessary for septin ring integrity during cytokinesis (Eluère et al., 2012; Kang et al., 2013; Wloka et al., 2011) and play distinct roles during the transition of septin filaments from an hourglass to a double ring (Chen et al., 2020). Bud3 also plays a regulatory role in Cdc42 polarization as a GDP-GTP exchange factor (GEF) for Cdc42 (Kang et al., 2014).

How are septins recruited to an incipient bud site? Cdc42 polarization occurs stepwise upon its activation by its GEFs, Bud3 and Cdc24, during the two temporal steps (T1 and T2) in the G1 phase (Kang et al., 2014). This biphasic Cdc42 polarization is likely linked to the recruitment and assembly of a septin ring. Notably, Cdc42 polarization during T1 is necessary for the axial landmark assembly and septin recruitment in **a** and α cells (Kang et al., 2018). Gic1 and Gic2, two related Cdc42 effectors, are involved in septin recruitment (Iwase et al., 2006; Sadian et al., 2013). However, overexpression of *CDC42* suppresses the lethality of cells lacking Rsr1 and both Gics and rescues the defects in septin recruitment (Kang et al., 2018), suggesting that Cdc42 can recruit septins directly or indirectly via a protein other than Gic proteins.

Previous genetic studies suggested that Axl2 is involved in septin organization. *AXL2* was identified as a dosage suppressor of the lethality of *spa2*Δ *cdc10-10,* a mutant defective in Cdc10, and Spa2, a polarisome component (Roemer et al., 1996). Overexpression of *AXL2* also suppresses temperature-sensitive (ts) growth of *gic1 gic2* (Gandhi et al., 2006) and a *cdc42* mutant defective in the septin ring assembly (Gao et al., 2007). Although it was reported that Axl2 interacts with Cdc42-GDP and Bud4 even in the absence of Bud3 (Gao et al., 2007), we found that Axl2 fails to associate with Bud4 in cells lacking Bud3 but expressing all other proteins at endogenous levels (Kang et al., 2012). This apparent inconsistency and other outstanding questions led us to investigate further the function of Axl2 in axial budding. Here, by combining live-cell imaging with genetic analyses, we found that Axl2 interacts with Bud3, Cdc42-GTP, and Cdc10. We also identified a *cdc42* mutation that disrupts its interaction with Axl2 and causes compromised septin recruitment. Our findings suggest that Axl2 promotes efficient septin recruitment to the axial bud site, while Axl2 may also function in septin recruitment in non-axially budding cells.

## RESULTS AND DISCUSSION

### Axl2 interacts with Bud3 and Cdc42-GTP

To test whether Axl2 interacts with Bud3, we performed pull-down assays using Axl2-TAP in wild-type (WT) or mutant strains expressing Bud3-Myc or Bud3ΔN-Myc (which lacks the N-terminal 259 residues). Bud3-Myc, but not Bud3ΔN-Myc, was associated with Axl2-TAP (Fig. 1A). This Axl2-Bud3 association was diminished by about 50% in an *axl1* mutant and almost completely abolished in a *bud4*Δ mutant. Since Axl2 fails to interact with Bud4 in *bud3*Δ cells (Kang et al., 2012), these results indicate that Axl2 associates with the Bud3-Bud4 complex and support the idea of the stepwise assembly of the axial landmark: first, the interaction between Bud4 and Bud3, and then the association of Axl2 and Axl1 with the Bud4-Bud3 dimer.

**Fig. 1.**
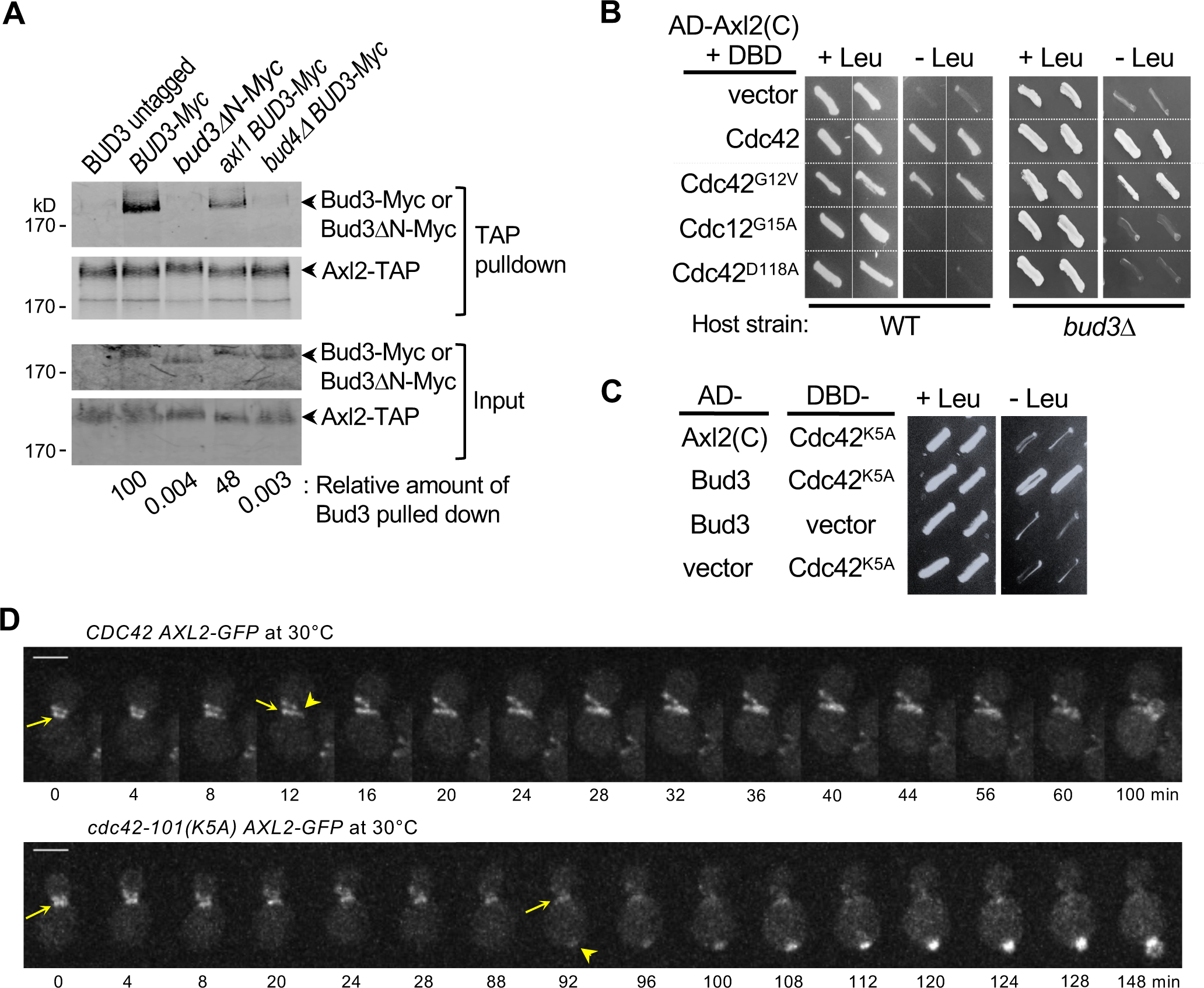
Axl2 interacts with Bud3 and Cdc42-GTP. **A.** Axl2-TAP pull-down assays using WT, *BUD3-Myc* (or *bud3ΔN-Myc*) mutants, and an untagged strain. **B.** Two-hybrid assays with AD-Axl2(C) and DBD fusions of WT or mutant Cdc42 in two host strains, as marked. **C.** Two-hybrid assays between DBD-Cdc42^K5A^ and AD-Axl2(C) or AD-Bud3. **D.** Time-lapse images of Axl2-GFP in WT and *cdc42-101* cells at 30°C. Axl2-GFP signals at the division site and incipient bud site in mother cells are marked with arrows and arrowheads, respectively. Size bars, 3 µm.

Is such ordered assembly of the axial landmark functionally significant? Since Bud3 is a Cdc42 GEF (Kang et al., 2014), we considered two possible scenarios. First, Bud3 may become an active Cdc42 GEF upon its interaction with Axl2. If so, Axl2 may associate with Cdc42-GDP, mediated by Bud3. Second, upon activation by Bud3, Cdc42-GTP may interact with Axl2 and bring it for the axial landmark assembly. To distinguish these, we tested the interaction between Axl2 and Cdc42 by a yeast two-hybrid assay. Expression of the *LEU2* reporter indicated that Axl2 interacts with WT Cdc42 and Cdc42^G12V^, a GTP-locked state in vivo. In contrast, Axl2 failed to interact with Cdc42^G15A^ and Cdc42^D118A^, which are in a GDP-locked or nucleotide-free state (Daubon et al., 2011; Davis et al., 1998) (Fig. 1B). These results suggest that Axl2 interacts with Cdc42-GTP. These findings also imply that Axl2 may not be necessary for Bud3 GEF activity and that Bud3 may not mediate the Axl2-Cdc42 interaction. Indeed, when we examined the interaction between Axl2 and Cdc42 in a *bud3*Δ mutant by a two-hybrid assay (Fig. 1B) and Axl2-TAP pull-down assays (Fig. S1), Axl2 interacted with Cdc42 similarly in WT and *bud3*Δ strains. Collectively, these data suggest that Axl2 interacts with Bud3 and with Cdc42-GTP independently of Bud3.

### *cdc42-101* is defective in the interaction with Axl2 and septin recruitment

To determine the functional significance of the interaction between Axl2 and Cdc42-GTP, we looked for a *cdc42* allele that encodes Cdc42 defective in interaction with Axl2. Among the *cdc42^ts^* alleles that arrest in G1 at 37°C (Kozminski et al., 2000), *cdc42-101^K5A^* is specifically defective in the axial budding pattern at 30°C, a semi-permissive temperature (Kang et al., 2014). Interestingly, Cdc42^K5A^ interacted normally with Bud3 but poorly with Axl2 in a two-hybrid assay (Fig. 1C). We then examined how this *cdc42* mutation impacts the localization of Axl2 at 30°C by time-lapse imaging. In WT cells, Axl2-GFP localized to the bud neck, forming a double ring, and in a patch next to the ring soon after cell division (93%, n = 36). In *cdc42-101* cells, however, Axl2-GFP frequently appeared as a patch at the pole distal to the cell division site (60%, n = 48; Fig. 1D). Despite the presence of all axial landmark proteins, this single residue substitution in Cdc42 seems to cause its poor interaction with Axl2, resulting in inefficient recruitment of Axl2 to the axial bud site. This also leads to the poor association of Axl1 with Bud4 in *cdc42-101* since the assembly of Axl2 with Bud3-Bud4 is necessary for the subsequent interaction of Axl1 with the complex (Kang et al., 2012; Kang et al., 2014). These findings suggest that Cdc42-GTP promotes the assembly of the axial landmark by bringing Axl2 near the cell division site.

Since *axl2*Δ mutants are viable, the lethality of *cdc42-101^K5A^* at 37°C suggests additional defects that may be related to (or distinct from) its poor interaction with Axl2. Interestingly, temperature-sensitive growth of *cdc42-101* was suppressed uniquely by overexpression of Gic1 among Cdc42 effectors (Kozminski et al., 2000). Based on this genetic interaction and Gic1’s role in septin recruitment (Iwase et al., 2006; Sadian et al., 2013), we postulated that *cdc42-101* might be defective in septin recruitment. To test this idea, we examined the localization of septin Cdc3-GFP in *cdc42-101* cells by time-lapse imaging after temperature upshift to 37°C. Septin recruitment occurred within 16 min after cytokinesis in WT mother cells (90%; n = 22). In contrast, less than 40% of the *cdc42-101* mother cells (n = 24) showed septin recruitment at 1 hr after cytokinesis, and even in those cells, new septins often failed to develop into a ring at 37°C (Figs. 2A & S2). We next examined the localization of Cdc3-GFP at 30°C, together with PBD (p21-binding domain)-RFP, a marker for Cdc42-GTP (Okada et al., 2013). New septins appeared next to the old septin ring at the division site in WT cells (95%, n = 60) but frequently at the pole distal to the division site in *cdc42-101* cells (59%, n = 68) (Fig. 2B). Collectively, these observations suggest that Cdc42^K5A^, which poorly interacts with Axl2, is defective in the recruitment of septins to the axial bud site at 30°C and more severely defective in septin recruitment at 37°C or above.

**Fig. 2.**
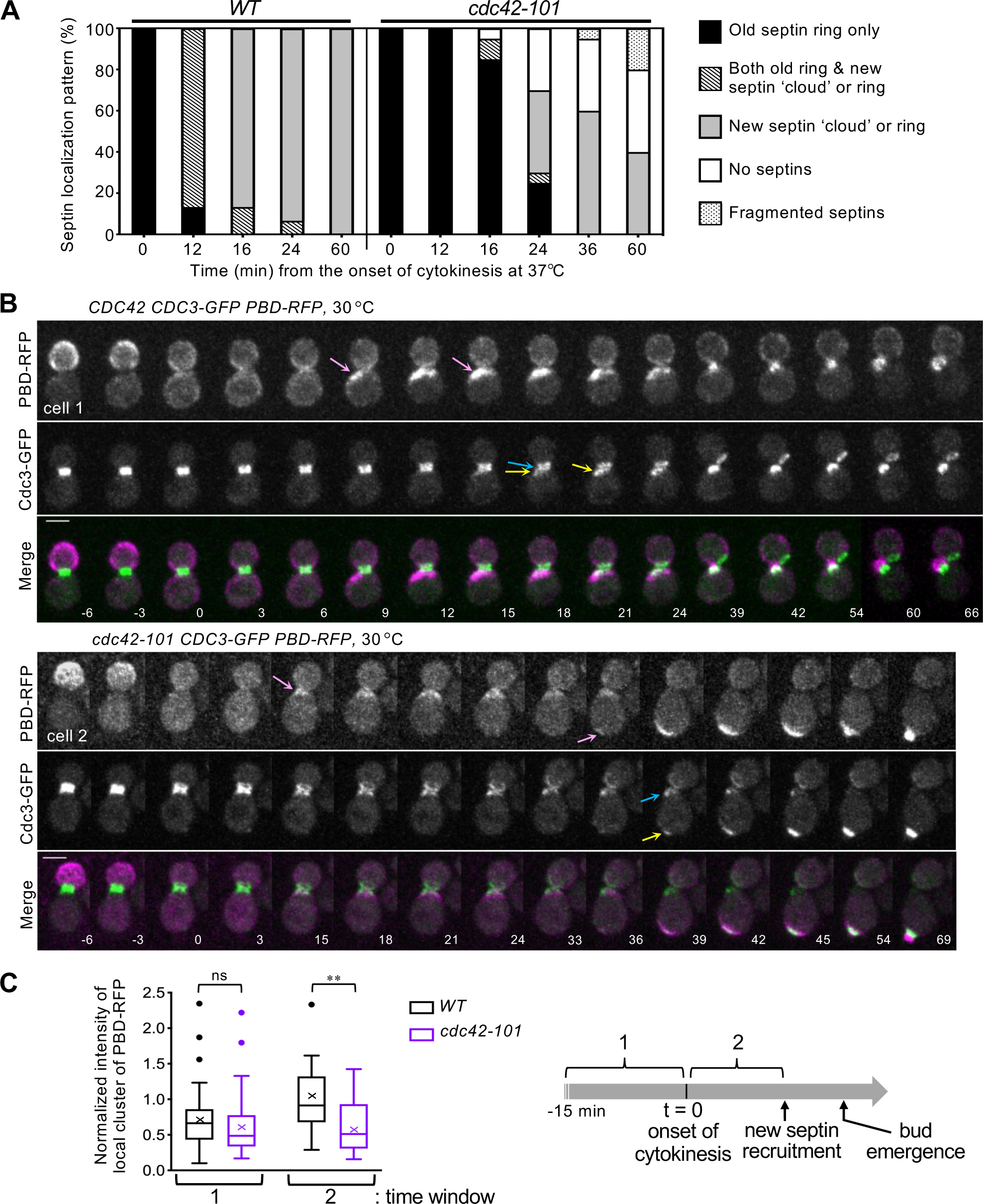
*cdc42-101^K5A^* is defective in septin recruitment. **A.** Cdc3-GFP localization patterns in WT and *cdc42-101* cells at 37°C, analyzed from time-lapse images (see **Fig. S2**). **B.** Time-lapse images of WT (cell 1) and *cdc42-101* (cell 2) expressing Cdc3-GFP and PBD-RFP at 30°C. The Cdc42-GTP clusters first appeared in early G1 and later at the incipient bud site in mother cells are marked with pink arrows; and old septin ring and new septins are marked with blue and yellow arrows, respectively. Numbers indicate time (in min) relative to the onset of cytokinesis (t = 0). Size bars, 3 µm. **C.** Fluorescence intensity of the PBD-RFP cluster in WT and *cdc42-101* cells at 30°C is normalized against the average peak values of WT in early G1 (the time window 2). The highest levels of PBD-RFP clusters in each time window are compared from the analyses of time-lapse images (n = 33, each strain) (see Materials and Methods).

In both WT and *cdc42-101* cells, new septins started to appear soon after the development of the first wave of Cdc42-GTP polarization in G1 at 30°C. We quantified the PBD-RFP cluster from the time-lapse images and compared the peak intensities in two time windows: 15 min before cytokinesis; and from the onset of cytokinesis to the first appearance of new septins (see Materials and Methods). The PBD-RFP level was similarly high in WT and *cdc42-101* cells before cytokinesis, indicating that Cdc42^K5A^ is not defective in its interaction with Cdc42 effectors (that contain the PBD) or with its GEFs. However, the PBD-RFP cluster level in early G1 was lower in *cdc42-101* than in WT (Fig. 2C), likely because of the compromised positive feedback loop of Cdc42 polarization due to inefficient coupling of the axial landmark (lacking Axl2) to the Rsr1 module (Kang et al., 2014; Lee et al., 2015). This defect is likely to result in inefficient septin recruitment to an axial bud site at 30°C.

### Axl2 interacts with Cdc10

Since Axl2 interacts with Cdc42-GTP but poorly with Cdc42^K5A^, which is compromised in septin recruitment, we hypothesized that Cdc42-GTP promotes septin recruitment via Axl2. To test this idea, we determined whether Axl2 interacts with any septin subunit. A two-hybrid assay suggested that Axl2 interacts with Cdc10 among the septin subunits Cdc3, Cdc10, Cdc11, and Cdc12 (Fig. 3A). We next examined the Axl2-Cdc10 interaction by a bimolecular fluorescence complementation (BiFC) assay. Expression of the N- and C-terminal fragments of Venus (V_N_ and V_C_) fused to Axl2 and Cdc10, respectively, resulted in YFP signals at the mother-bud neck and the division site (Fig. 3B). In contrast, co-expression of Axl2-V_N_ and Cdc11-V_C_ (or Cdc12-V_C_) did not show any YFP signals. Unlike other septin subunits, Cdc10 does not have a C-terminal extension (CTE). The proximity of V_C_ (at the C terminus of septins) to V_N_ might potentially result in more efficient recovery of YFP with Cdc10-V_C_ compared to other septin-V_C_ fusions even when Axl2-V_N_ interacted with a common domain in septin subunits. To test this possibility, we expressed Cdc11ΔC-V_C_, which lacks its CTE (a.a. 357–415). Expression of *cdc11ΔC*-*V_C_* (as a sole copy of *CDC11*) did not cause any growth defect, consistent with a previous report that the Cdc11 CTE is dispensable for the septin assembly (Versele et al., 2004). Notably, co-expression of Cdc11ΔC-V_C_ and Axl2-V_N_ also did not recover any YFP signal (Fig. 3B). Collectively, these data suggest that Axl2 interacts closely with Cdc10.

**Fig. 3.**
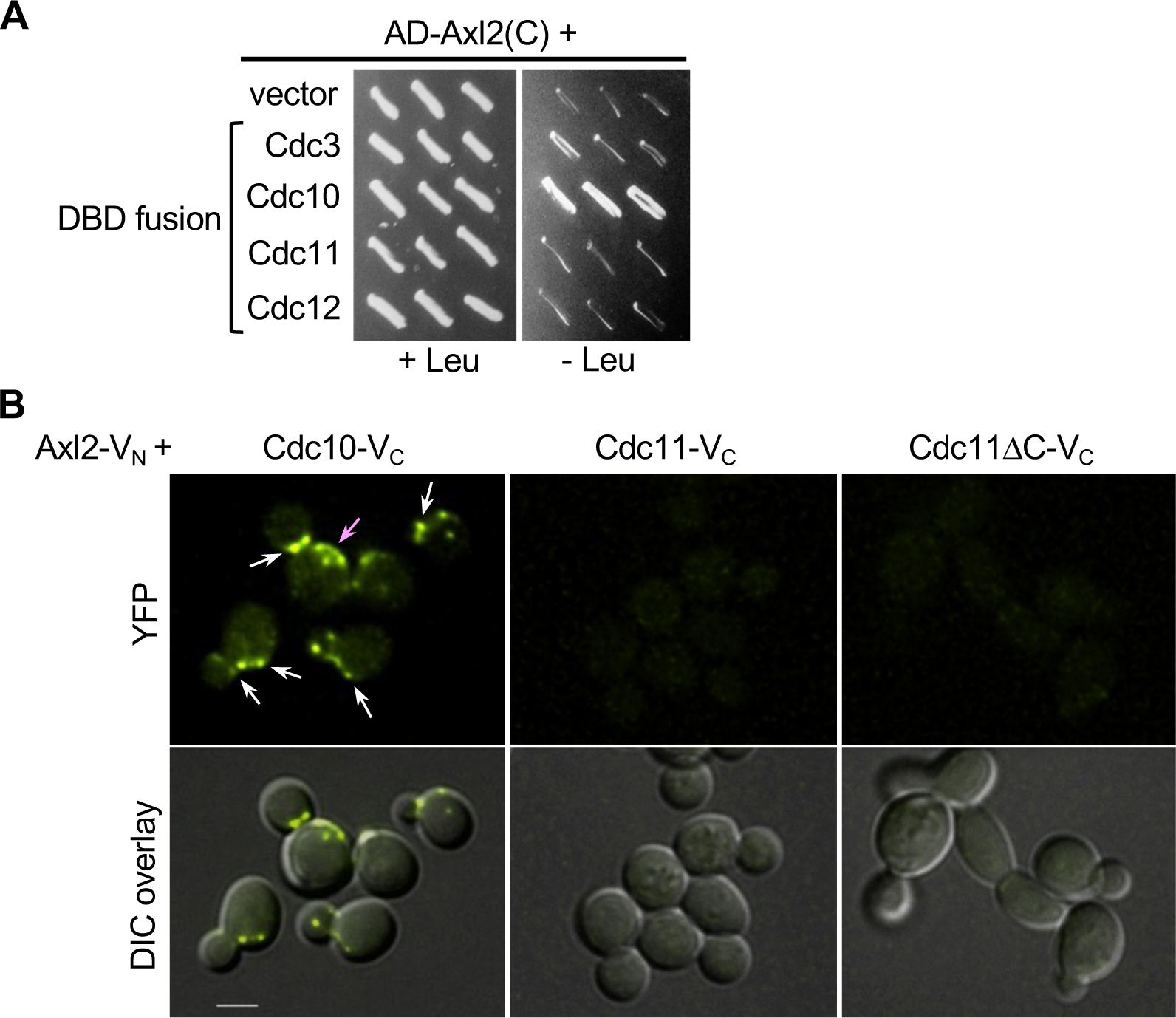
Axl2 interacts with Cdc10. **A.** Two-hybrid assays between AD-Axl2(C) and DBD-septin subunits (or vector control). **B.** BiFC assays between Axl2-V_N_ and Cdc10-V_C_, Cdc11-V_C_, or Cdc11ΔC-V_C_. YFP signals are marked at the bud neck and division site (white arrows) and in patches often clustered near the division site (pink arrow). Size bars, 3 µm.

### Axl2 may facilitate septin recruitment by interacting with Cdc10

What is the functional significance of the Axl2 - Cdc10 interaction? The Cdc10 dimer is the core of the septin protofilaments and associates with one of two trimers, containing either Cdc11 or Shs1, to form an octamer (Weems and McMurray, 2017). We hypothesized that Axl2 might recruit septins by interacting with Cdc10 at the incipient bud site where Cdc42 is polarized. To test this idea, we first examined how Axl2 affects the localization of septins. Cdc10-GFP, which displayed no noticeable defects in WT, frequently mislocalized in *axl2*Δ cells, appearing diffused in the cytoplasm or at the bud tip (Fig. 4A). In contrast, Cdc3-GFP or Cdc11-tdTomato localized similarly in WT and *axl2*Δ cells (Fig. S3, A & B). These observations suggest that Axl2 interacts mainly with Cdc10, although GFP tagging of Cdc10 might have a minor impact on its function (see below).

**Fig. 4.**
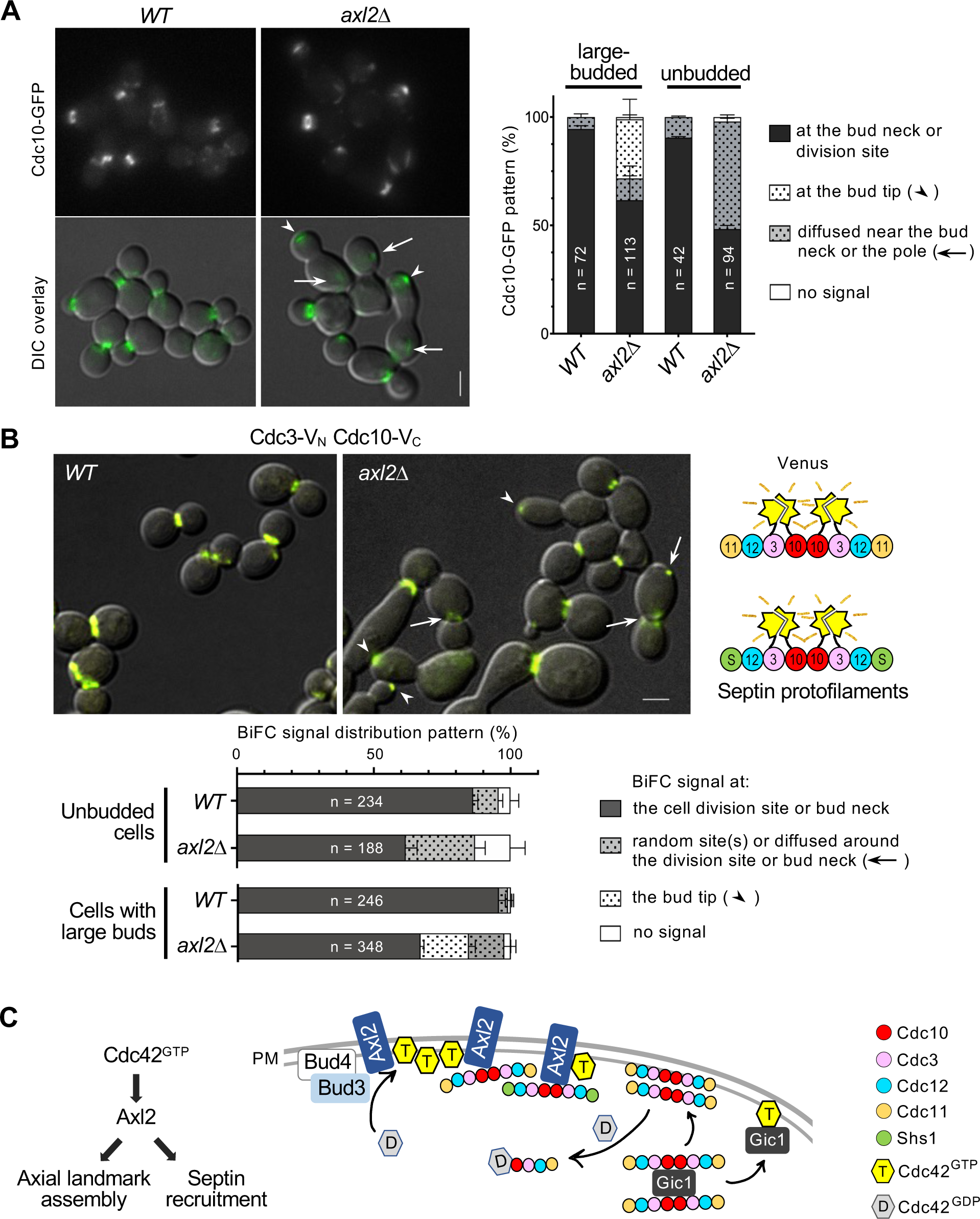
Abnormal septin recruitment in *axl2*Δ cells. **A.** Localization of Cdc10-GFP in WT and *axl2*Δ cells at ∼24°C, and the analysis of the localization patterns in unbudded or large-budded cells. Size bars, 3 µm. **B.** BiFC assays between Cdc3-V_N_ and Cdc10-V_C_ in WT and *axl2*Δ cells. The YFP localization patterns are analyzed from the total number (n) of cells, as marked, pulled from three imaging sets. The scheme on the right shows Venus reconstitution expected in septin protofilaments. **C.** Model for Cdc42 coupling the axial landmark assembly and septin recruitment via Axl2. For simplicity, septins are shown only as octamers (see text).

Next, we performed BiFC assays to test whether Axl2 affects the interaction between Cdc10 and Cdc3. Expression of Cdc3-V_N_ and Cdc10-V_C_ showed YFP signals at the bud neck and cell division site in WT cells but additionally at the bud tip in *axl2*Δ cells (Fig. 4B). The YFP signals at the division site in *axl2*Δ cells often appeared as a less tightly organized ring (see Materials and Methods). These observations suggested that Axl2 functions in the efficient recruitment of septins but also raised a question of whether some molecules of Cdc3 mislocalize in *axl2*Δ cells at least transiently. Indeed, Cdc3-mCherry mislocalized to the bud tip in some *axl2*Δ cells expressing Cdc10-GFP, even though no such mislocalization of Cdc3-GFP was observed in *axl2*Δ cells (carrying untagged *CDC10)* (see Fig. S3A). GFP (or V_C_) tagging of Cdc10 is thus likely to cause a subtle defect that is exacerbated in the absence of Axl2, as also indicated by a small percentage of these *axl2*Δ cells with abnormal cell shapes. We also examined the homotypic interaction of Cdc10 by BiFC assays using a diploid strain expressing Cdc10-V_N_ and Cdc10-V_C_. YFP signals appeared at the bud tip and also diffused at the division site in some *axl2*Δ cells (Fig. S4). Collectively, these results suggest that Axl2 is involved in the efficient recruitment of septins to the cell division site in both haploid and diploid cells. While this is consistent with similar localization patterns of Axl2 to the bud neck and the division site in all cell types, it is yet to be determined whether Axl2 plays any role in septin organization in diploid cells.

### Limitations of the study and a model

Our findings in this study suggest that Axl2 couples septin recruitment to the axial landmark assembly by interacting with Bud3, Cdc42-GTP, and Cdc10. Yet several questions remain, including whether these interactions are direct or whether these interactions occur sequentially or simultaneously. It also remains an open question whether Axl2 interacts with Cdc10 as a monomer, dimer, or within septin protofilaments. Axl2 is a transmembrane protein, whose glycosylation is critical for its function (Powers and Barlowe, 1998; Roemer et al., 1996; Sanders et al., 1999), but direct evidence is limited regarding when and where Axl2 interacts with Cdc42 or Cdc10.

Despite these limitations, we note that Axl2 shares similar binding partners with Gic1, a yeast analog of the mammalian Borg proteins involved in the septin organization (Farrugia and Calvo, 2016; Iwase et al., 2006; Sadian et al., 2013). Gic1 binds to Cdc42-GTP and Cdc10 and scaffolds septin filaments into long, flexible cables, while Cdc42-GDP binds to Cdc10 and dissociates septin filaments (Sadian et al., 2013). Axl2 may share a partially redundant role with Gic1 in septin recruitment, although the mechanisms of their actions are likely different (Fig. 4C). In **a** or α cells, this role of Axl2 is likely linked to the axial landmark via its interaction with the Bud3-Bud4 complex. After the joining of Axl2 and Axl1, this complex becomes the functional axial landmark that interacts with the Rsr1 GTPase module (Kang et al., 2012). Since Axl2 can interact with Cdc42 in *bud3*Δ cells (this study), Axl2 may also function in septin recruitment independently of the axial landmark, as previously suggested (Gao et al., 2007). However, this is also likely dependent on Axl2’s interaction with Cdc42-GTP, which is polarized by a default mechanism in the absence of a spatial cue (Kang et al., 2018). Remarkably, a *cdc42* mutant defective in interaction with Axl2 displays abnormal septin recruitment (this study) and is suppressed uniquely by overexpression of Gic1 among Cdc42 effectors (Kozminski et al., 2000). These data further support the functional link between Axl2 and Gic1. We thus propose that Cdc42 orchestrates both axial landmark assembly and septin recruitment via Axl2 in haploid cells. Further studies are required to fully understand these interactions and to delineate the mechanisms of septin recruitment to the incipient bud site.

## MATERIALS AND METHODS

### Strains, plasmids, and growth conditions

Standard methods of yeast genetics, DNA manipulation, and growth conditions were used (Guthrie and Fink, 1991). Yeast strains used for imaging express tagged proteins under their native promoters from the chromosomes. Yeast strains were grown in rich yeast medium YPD (yeast extract, peptone, dextrose) or synthetic complete (SC) containing 2% dextrose as a carbon source unless stated otherwise. The strains and plasmids used in this study are listed in supplementary material Tables S1 and S2, and brief descriptions of construction methods are provided below the Tables.

### Microscopy and image analysis

Cells were grown in SC medium (with 2% dextrose) overnight and then freshly subcultured for 3 – 4 h before mounting on a slab containing the same medium and 2% agarose. Time-lapse imaging was performed essentially as previously described (Kang et al., 2014; Lee et al., 2015) using a spinning-disk confocal microscope (Ultra-VIEW VoX CSU-X1 system; PerkinElmer) equipped with a 100×, 1.45 NA Plan Apochromat objective lens (Nikon), 440-, 488-, 515-, and 561-nm solid-state lasers (Modular Laser System 2.0; PerkinElmer), and a back-thinned electron-multiplying CCD (charge-coupled device) camera (ImagEM C9100-13; Hamamatsu Photonics) on an inverted microscope (Ti-E; Nikon). Images were captured (9 Z stacks, 0.3 µm Z steps) every 3 or 4 min at 30°C or after shifting to 37°C (as indicated), and maximum intensity projections of Z stacks were used to make Figs. 1D, 2, and S2. Throughout the paper, numbers in time-lapse images indicate time (in min) from the onset of cytokinesis (t = 0), which was identified based on the splitting of the septin ring, and size bars indicate 3 µm.

To analyze Cdc42 polarization, the PBD-RFP clusters were quantified by a threshold method using an ImageJ ((National Institutes of Health) macro, as previously described (Okada et al., 2013; Okada et al., 2017). Briefly, average intensity projection images were generated from five best-focused Z stacks, and then a threshold method was used after background subtraction to quantify the PBD-RFP clusters in mother cells at each time point. The peak PBD-RFP level was identified in each period: 1) from t = −15 min to t = 0 (the onset of cytokinesis) or 2) from the onset of cytokinesis until the appearance of new septins. These peak values were plotted after normalizing against the average of the WT peak values during the second time window (i.e., early G1) in Fig. 2C.

To compare the localization of septin subunits in WT and *axl2*Δ cells, cells were grown as described above. Slides were prepared on an agarose slab as described above, and static images were captured under the same conditions (13 Z stacks, 0.3 µm Z steps) at ∼24°C using an inverted widefield fluorescence microscope (Ti-E; Nikon) equipped with a 100x/1.45 NA Plan Apochromat Lambda oil immersion objective lens, YFP, FITC/GFP, and mCherry/TexasRed filters from Chroma Technology, DIC optics, an EM CCD (Andor iXon Ultra 888) (Andor Technology), and the software Nis elements (Nikon). Maximum intensity projections of Z stacks (without deconvolution) were used to make Figs. 4A and S3.

### BiFC assays

BiFC assay is based on the fluorescence recovery by the interaction of two proteins, each of which is fused to the N- or C-terminal fragment of Venus (V_N_ or V_C_) (Kerppola, 2006). Yeast strains expressing a combination of test proteins fused to V_N_ or V_C_ were grown, and slides were prepared with an agarose slab (as described above). Images were captured (5 Z stacks, 0.4 µm Z steps) at ∼24°C using an inverted widefield fluorescence microscope (Ti-E; Nikon) and the YFP filter (see above). Localization patterns of BiFC signals were analyzed by applying the same threshold to all images (to highlight the localized YFP signals) after subtracting background using the ‘rolling ball’ method in ImageJ or Fiji software. Abnormal patterns of BiFC signals (such as a less tightly organized ring) in *axl2*Δ cells were more clearly identified by highlighting each pixel above a threshold, and the same threshold was applied to compare WT and *axl2*Δ cells. These patterns were analyzed in unbudded cells or cells with large buds from three independent image sets, excluding a small percentage of *axl2*Δ cells that had abnormal shapes or a cytokinesis defect. To make figures, YFP images were deconvolved by the Iterative Constrained Richardson-Lucy algorithm (Nis Elements), and a single best-focused Z slice was overlaid with DIC (Figs. 3B, 4B, and S4).

### In vitro binding assays and immunoblotting

All tagged proteins were expressed in yeast strains at their chromosomal loci (see Tables S1), except GST-Cdc42. These strains were grown to the mid-log phase (OD_600_ ∼ 1.0) in YPD at 30°C. GST or GST-Cdc42 was expressed from a plasmid (pRD56 or pEGKT-CDC42) (Gao et al., 2007; Park et al., 1993), and strains carrying these plasmids were initially grown in SC (with 2% sucrose) lacking Ura and then for additional 5 hrs after adding 2% galactose. TAP pull-down assays were carried out at 4°C, as previously described (Kang et al., 2014). Briefly, about 100 OD_600_ units of cells were used to prepare cell lysates using a lysis buffer (50 mM HEPES, pH 7.6, 300 mM KCl, 1 mM EGTA, 1 mM MgCl_2,_ 10% glycerol, and 1% Triton X-100) with a cocktail of protease inhibitors. The crude cell lysates were centrifuged for 10 min at 10,000 X g, and the supernatant (S10 fraction) was used for subsequent assays. For TAP pull-down assays, the S10 fraction was incubated with 25 µl of IgG-Sepharose (Cytiva, Marlborough, MA) for 1 h at 4°C by rocking. After washing the beads with the same lysis buffer, proteins were eluted from the beads and then subjected to immunoblotting. To determine the Cdc42-Axl2 interaction by TAP pull-down assays, cell lysates were prepared from ∼ 100 OD_600_ units of cells expressing GST or GST-CDC42 using a lysis buffer (50 mM HEPES, pH 7.6, 100 mM KCl, 1 mM EGTA, 1 mM MgCl_2,_ 10% glycerol, and 0.33% Triton X-100) with a cocktail of protease inhibitors. These cell lysates were then incubated with TAP-Axl2 bound IgG-Sepharose beads for 1 h at 4°C by rocking, followed by subsequent steps of washing, elution, and immunoblotting. Myc-, GST-, and TAP-tagged proteins were detected using anti-Myc antibody 9E10 (a gift of M. Bishop, University of California-San Francisco; used at a dilution of 1:1,000), rabbit anti-GST antibody (Santa Cruz Biotechnology, cat#sc-459, Santa Cruz, CA; used at a dilution of 1:500), and rabbit monoclonal anti-calmodulin binding protein (Upstate Cell Signaling Solutions, cat# 05-932, Temecula, CA; used at a dilution of 1: 5,000), respectively. Protein bands were then detected with Alexa Fluor ^R^ 680 goat anti-rabbit IgG (Invitrogen, cat# A32734; used at a dilution of 1:10,000) or IRDye^R^ 800CW conjugated goat anti-mouse IgG secondary antibodies (LI-COR Biosciences, cat# 926-32210, Lincoln, Nebraska; used at a dilution of 1:10,000) using the LI-COR Odyssey system (LI-COR Biosciences, Lincoln, Nebraska). These assays were repeated with two independent protein preparations, and the relative recovery of Bud3 compared to the input in each TAP pulldown assay was normalized against the same from the WT *BUD3-Myc* strain (Fig. 1A). The average percentage of GST-Cdc42 pulled-down relative to input was shown from two independent protein preparations (Fig. S1). A set of the original immunoblots is shown for blot transparency in Fig. S5.

### Yeast two-hybrid assay

Yeast two-hybrid assays were performed as previously described (Kang et al., 2014) by expressing activation domain (AD) fusions using pJG4-5 and DNA-binding domain (DBD) fusions using pEG202 (Gyuris et al., 1993). Two-hybrid assays were performed by patching 3 independent transformants of the host strain – *WT* (EGY48) or *bud3Δ* (HPY3657) – with a combination of AD and DBD plasmids on SGal plates lacking His and Trp (+ Leu) and SGal plates lacking His, Trp, and Leu (-Leu). Expression of the *LEU2* reporter was monitored based on the growth on SGal -Leu for 3 – 4 days at 30°C. The cytoplasmic domain (C) of Axl2 (aa 529 – 823) and Bud3 (aa 1-656), which carries the GEF domain, were fused to AD. Plasmids and strains used in two-hybrid assays are listed in Tables S1 and S2.

### Statistical analysis

Data analysis was performed using Prism 8 (GraphPad Software). Error bars in bar graphs (in Figs. 4 & S4) indicate SEM (standard error of the mean). In the box graph (Fig. 2C), quartiles and median values are shown together with the mean (marked with x). The Tukey method was used to create the whiskers, which indicate variability outside the upper and lower quartiles. Any point outside those whiskers (indicated as dots) is considered an outlier. A two-tailed student’s t-test was performed to determine statistical differences between two sets of data: ns (not significant) for p ≥ 0.05, *p < 0.05, and **p < 0.01.

## Abbreviations

GEF: GDP-GTP exchange factor
WT: wild type
BiFC: bimolecular fluorescence complementation
AD: activation domain
DBD: DNA-binding domain
TAP: tandem affinity purification
PBD: p21-binding domain
ts: temperature-sensitive

## ACKNOWLEDGEMENTS

We thank K. Kozminski, M. Peter, M. A. McMurray, E. Bi, and M. Farkasovsky for strains and plasmids. This work was supported by NIH grants (R01-GM114582 and R21-AG060028).

## SUPPLEMENTAL MATERIAL

Figures S1 – S5

Tables S1 – S2

## SUPPLEMENTAL FIG. LEGENDS

**Fig. S1.**
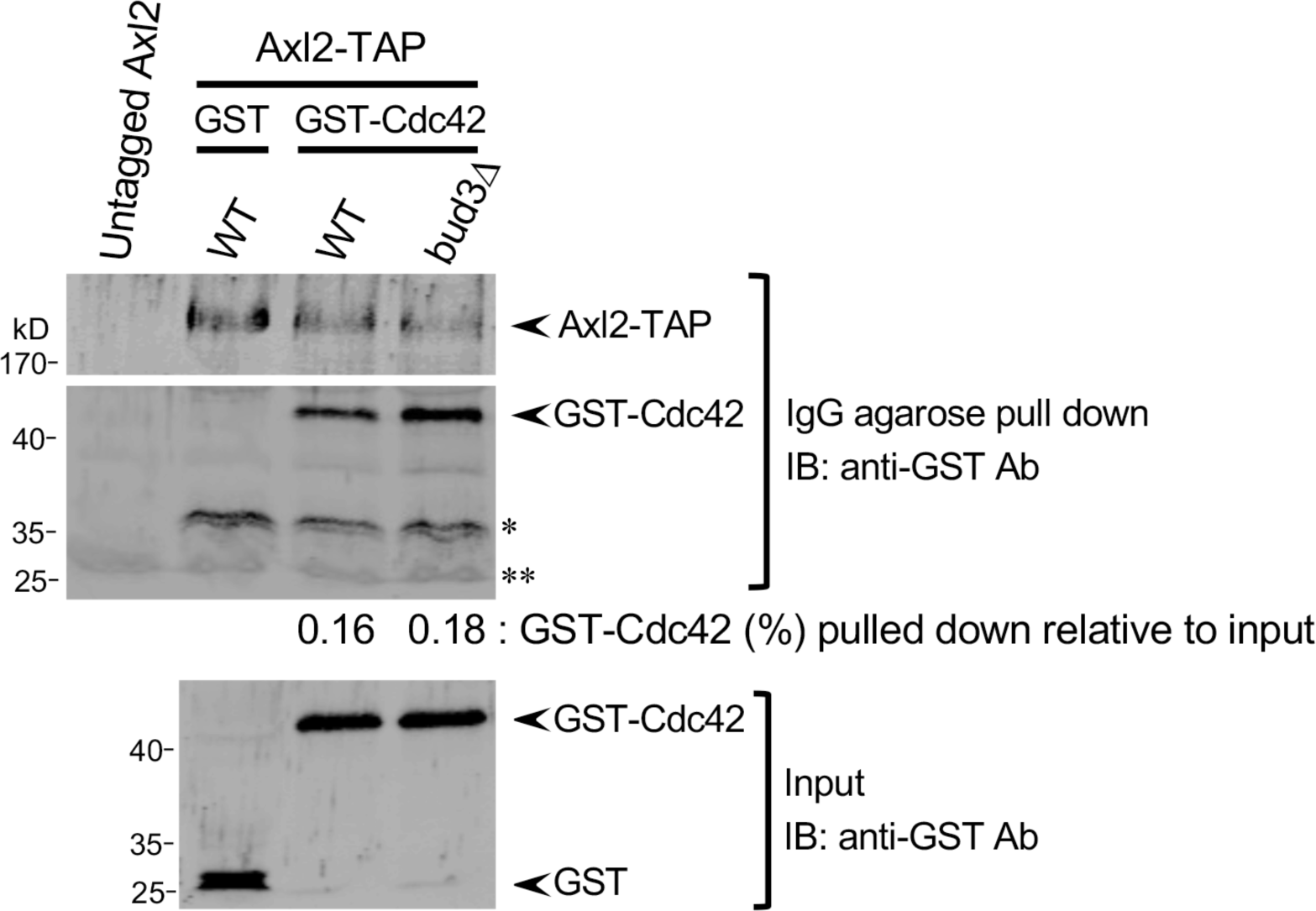
Interaction between Cdc42 and Axl2 by TAP-pull-down assays. Axl2-TAP was pulled down after incubating with GST-Cdc42 (or GST control) prepared from the *WT* and *bud3*Δ strains expressing Axl2-TAP. Lysates (input) and proteins pulled with IgG agarose were analyzed by immunoblotting with anti-GST antibodies, which also recognize the protein A tag in Axl2-TAP. GST-Cdc42 pull-down (%) shows the average amount recovered relative to input from two-independent protein preparations. The bands marked with * and ** are likely Axl2-TAP cleavage products and IgG light chain, respectively.

**Fig. S2.**
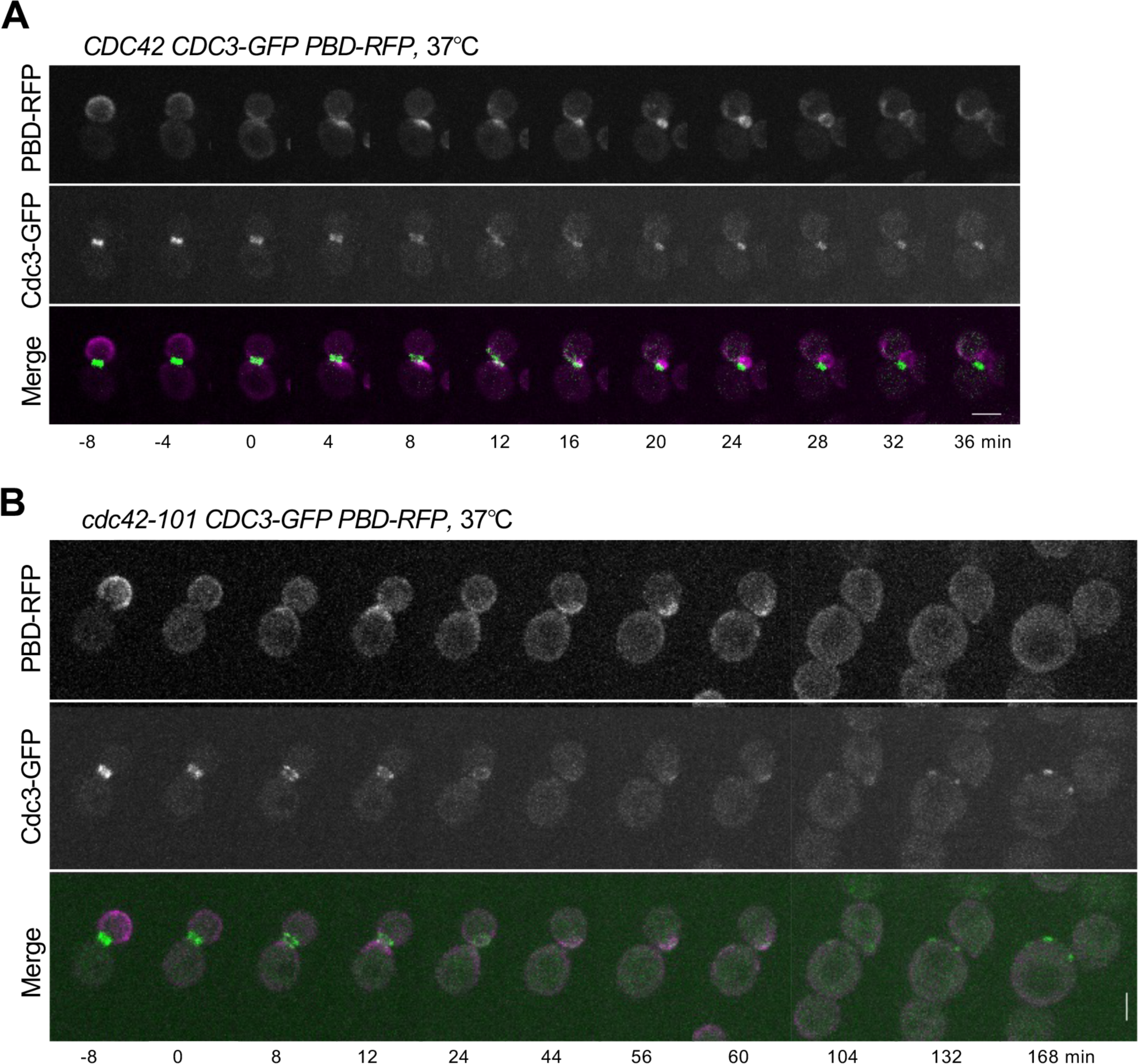
Time-lapse images of WT and *cdc42-101* cells expressing Cdc3-GFP and PBD-tdTomato at 37°C. Images were captured every 4 min, starting at 1 h after shifting to 37°C from ∼24°C. Images of WT (**A**) and *cdc42-101* (**B**) cells are shown at the selected time points. Time (min) is marked relative to the onset of cytokinesis (t = 0). Size bars, 3 µm.

**Fig. S3.**
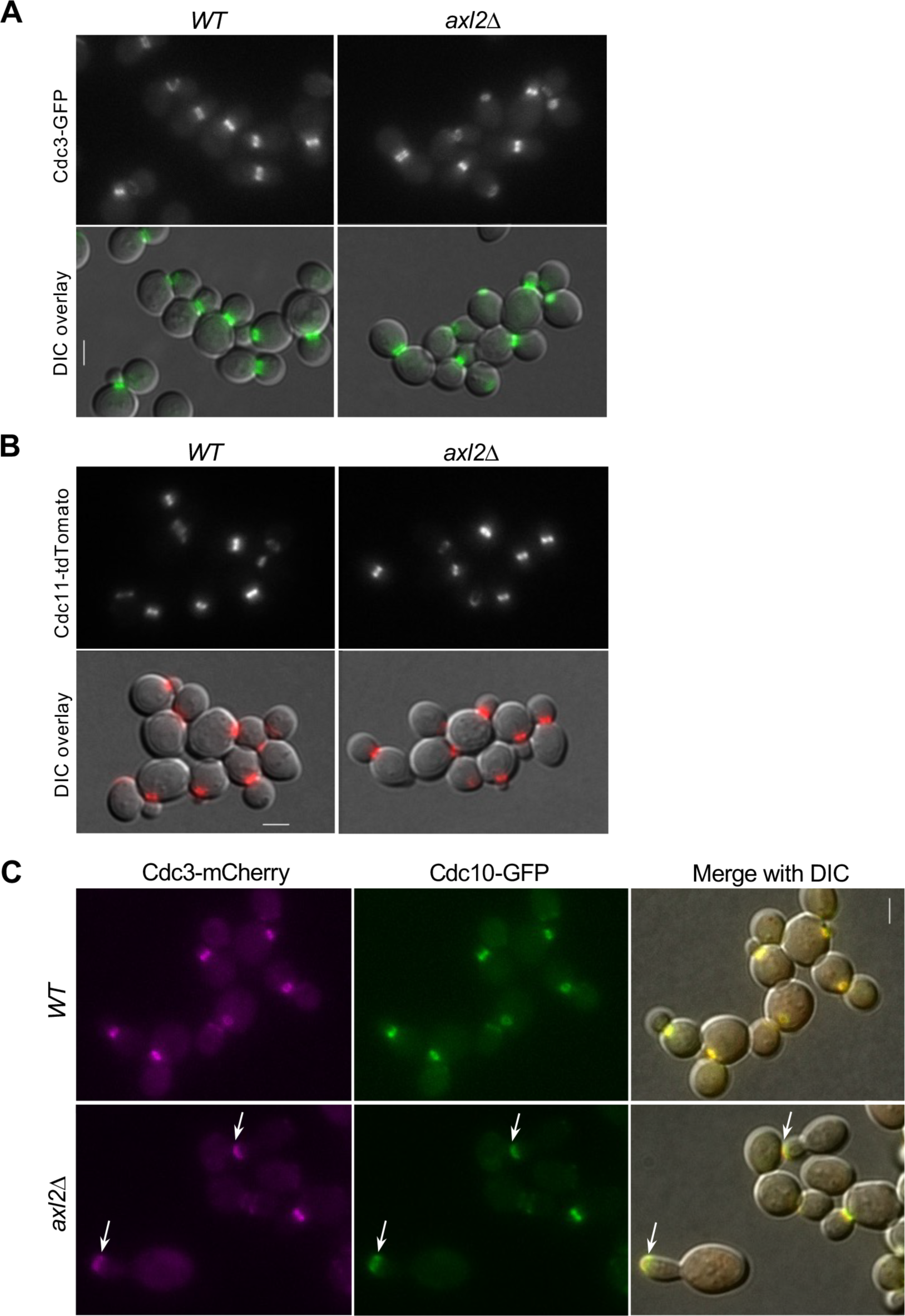
Localization of septins in WT and mutant cells at 24°C. **A.** Localization of Cdc3-GFP in WT and *axl2*Δ cells. Size bar, 3 µm. **B.** Localization of Cdc11-tdTomato in WT and *axl2*Δ cells. Size bar, 3 µm. **C.** Co-localization of Cdc3-mCherry and Cdc10-GFP in WT and *axl2*Δ cells. A small number of *axl2*Δ cells showed abnormal localization of both proteins to the bud tip (marked with arrows).

**Fig. S4.**
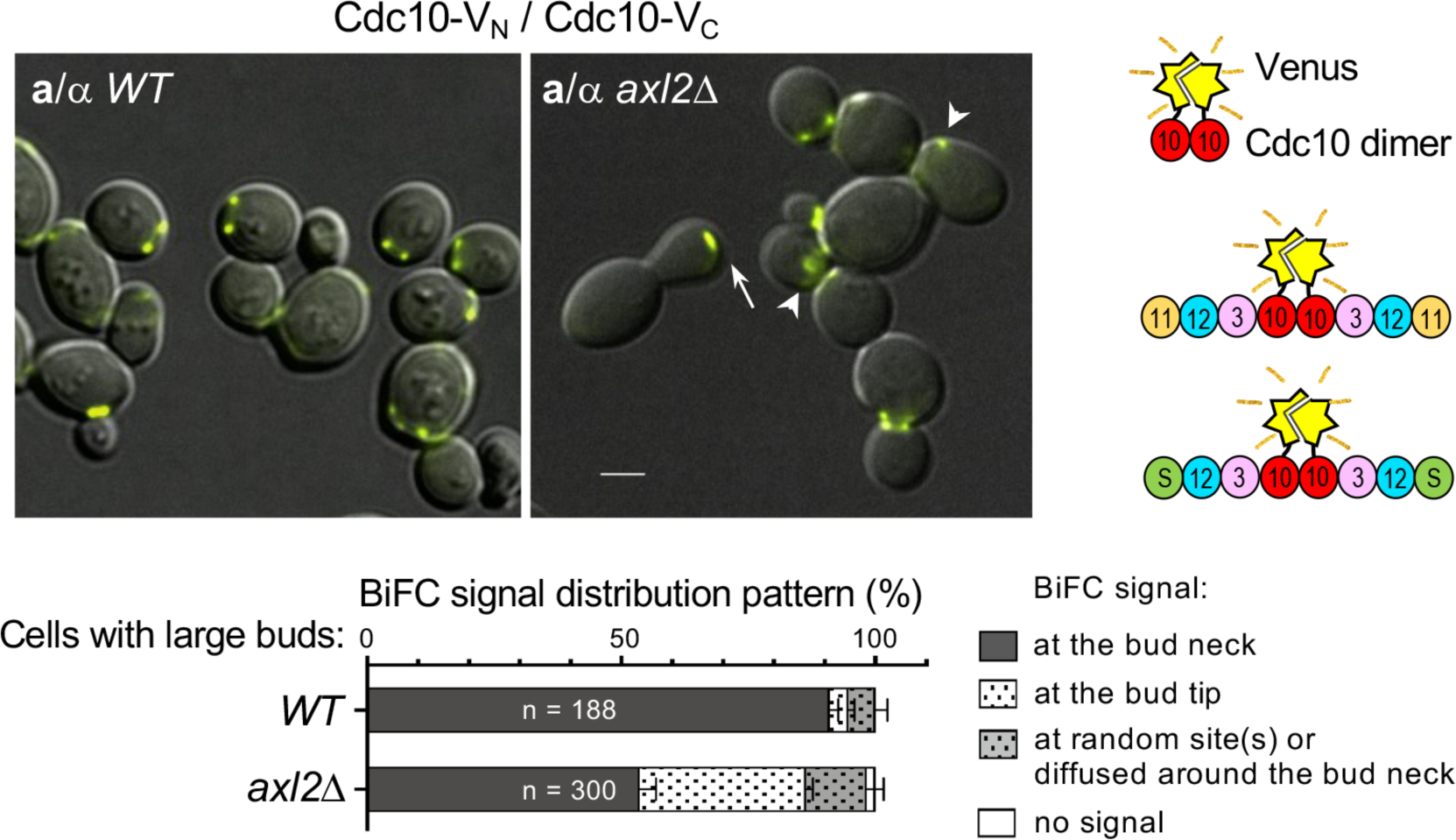
BiFC assays with Cdc10-V_N_ and Cdc10-V_C_ in diploid WT and *axl2*Δ cells. The abnormal appearance of YFP signals in *axl2*Δ cells is marked with an arrow (at the bud tip) or arrowheads (diffused ring at the cell division site). Size bar, 3µm. The BiFC signal distribution patterns are analyzed from the total number (n) of cells (pulled from three imaging sets). Schemes (right) depict the possible reconstitution of Venus from the Cdc10 homodimer or Cdc10 dimers within the septin protofilaments.

**Fig. S5.**
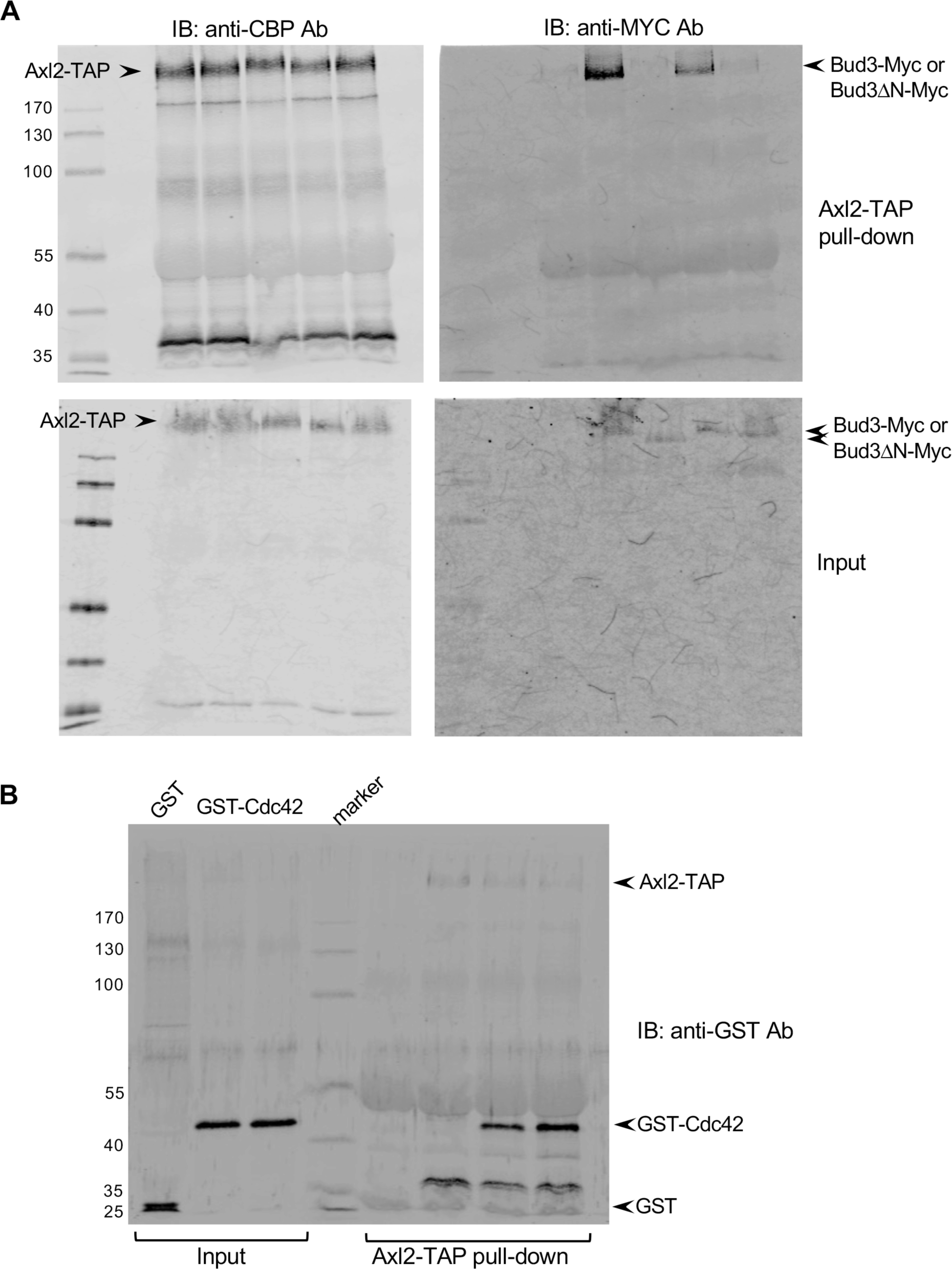
Blot transparency. **A.** Axl2-TAP was detected using anti-CBP antibodies (two blots on the left), and Bud3-Myc was detected with anti-Myc antibodies (two blots on the right). See Fig. 1A. **B.** GST-Cdc42 or GST was detected using anti-GST antibodies, which also recognize the protein A tag in Axl2-TAP. See Fig. S1.

**Supplementary Table S1.** Yeast strains used in this study.

| Strain <sup>a</sup> | Relevant Genotype | Source |
| --- | --- | --- |
| DDY1301<sup>@</sup> | *MATa CDC42-LEU2 his3-Δ200 leu2-3, 112 lys2-801 ura3-52* | (Kozminski et al., 2000) |
| DDY1304<sup>@</sup> | *MATa cdc42-101(K5A)-LEU2* | (Kozminski et al., 2000) |
| EGY48 | *MATα his3 trp1 ura3 LexA<sub>op(x6)</sub>-LEU2* | (Gyuris et al., 1993) |
| HPY3657 | *MATα his3 trp1 ura3 LexA<sub>op(x6)</sub>-LEU2 bud3Δ::URA3* | This study |
| YEF473A<sup>#</sup> | *MATa his3-Δ200 leu2-Δ1 lys2-801 trp1-Δ63 ura3-52* | (Bi and Pringle, 1996) |
| HPY16<sup>*</sup> | *MATa ura3-52 trp1-Δ63 leu2 his3-Δ1 pep4-3* | (Park et al., 1993) |
| BY4741<sup>@</sup> | *MATa his3Δ1 leu2Δ0 met15Δ0 ura3Δ0* | Open Biosystem |
| HPY1445<sup>@</sup> | *MATa AXL2-TAP-HIS3* | Open Biosystem |
| HPY1454<sup>@</sup> | *MATa bud3Δ::URA3 AXL2-TAP-HIS3* | This study |
| HPY2446<sup>*</sup> | *MATa bud3Δ::HIS3* | This study |
| HPY3130<sup>*</sup> | *MATa AXL2-TAP-HIS3* | This study |
| HPY3119<sup>*</sup> | *MATa BUD3-Myc<sub>13</sub>-KAN AXL2-TAP-HIS3* | This study |
| HPY3131<sup>*</sup> | *MATa bud3Δ::URA3 bud3ΔN(aa 2-260)-Myc<sub>13</sub>-KAN-LEU2 AXL2-TAP-HIS3* | This study |
| HPY3195<sup>*</sup> | *MATa bud4Δ::LEU2 BUD3-Myc<sub>13</sub>-KAN AXL2-TAP-HIS3* | This study |
| HPY3196<sup>*</sup> | *MATa axl1::URA3 BUD3-Myc<sub>13</sub>-KAN AXL2-TAP-HIS3* | This study |
| HPY3638<sup>@</sup> | *MATα CDC42-LEU2 AXL2-GFP-KAN* | This study |
| HPY3639<sup>@</sup> | *MATα cdc42-101(K5A)-LEU2 AXL2-GFP-KAN* | This study |
| HPY2580<sup>@</sup> | *MATa CDC42-LEU2 GIC2-PBD(W23A)-tdTomato-URA3 CDC3-GFP-LEU2* | This study |
| HPY2578<sup>@</sup> | *MATa cdc42-101(K5A)-LEU2 GIC2-PBD(W23A)- tdTomato-URA3 CDC3-GFP-LEU2* | This study |
| HPY3377<sup>#</sup> | *MATa CDC10-GFP-HIS3* | This study |
| HPY3419<sup>#</sup> | *MATa axl2Δ::KAN CDC10-GFP-HIS3* | This study |
| HPY3652<sup>#</sup> | *MATa AXL2-V<sub>N</sub>-HIS3 CDC10-V<sub>C</sub>-TRP1* | This study |
| HPY3641<sup>#</sup> | *MATa/MATα CDC10-V<sub>N</sub>-HIS3/CDC10-V<sub>C</sub>-TRP1* | This study |
| HPY3903<sup>#</sup> | *MATa/MATα CDC10-V<sub>N</sub>-HIS3/CDC10-V<sub>C</sub>-TRP1 axl2Δ::KAN/axl2Δ::KAN* | This study |
| HPY3844<sup>#</sup> | *MATα CDC3-V<sub>N</sub>-TRP1 CDC10-V<sub>C</sub>-HIS3* | This study <sup>b</sup> |
| HPY3908<sup>#</sup> | *MATa CDC3-V<sub>N</sub>-TRP1 CDC10-V<sub>C</sub>-HIS3 axl2Δ::KAN* | This study <sup>b</sup> |
| HPY3915<sup>#</sup> | *MATa AXL2-V<sub>N</sub>-HIS3 CDC11-V<sub>C</sub>-TRP1* | This study <sup>b</sup> |
| HPY3912<sup>#</sup> | *MATa AXL2-V<sub>N</sub>-HIS3 cdc11(Δ357–415)-V<sub>C</sub>-TRP1* | This study<sup>b</sup> |
| HPY3917<sup>#</sup> | *MATa CDC11-tdTomato-NAT* | This study |
| HPY3930<sup>#</sup> | *MATa CDC11-tdTomato-NAT axl2Δ::HIS3* | This study |
| HPY3935 | *MATa CDC3-GFP-LEU2* | This study |
| HPY3932<sup>#</sup> | *MATα CDC3-GFP-LEU2 axl2Δ::HIS3* | This study |
| HPY3727<sup>#</sup> | *MATa CDC3-mCherry-LEU2 CDC10-GFP-HIS3* | This study |
| HPY3729<sup>#</sup> | *MATa axl2Δ::KAN CDC3-mCherry-LEU2 CDC10-GFP-HIS3* | This study |

<sup>a</sup> Strains marked with <sup>#</sup> are congenic to YEF47A (Bi and Pringle, 1996); strains marked with @ are derived from DDY1301 or DDY1304 (in S288C background); and strains marked with \* are isogenic to HPY16 (Park et al., 1993), except as indicated.
<sup>b</sup> Derived using YO802 (*CDC12-V<sub>C</sub>::KanMX6*), YO685 (*CDC3-V<sub>C</sub>::HIS3MX6*), YEF5692 (*CDC12-V<sub>N</sub>::TRP1*), and YEF5689 (*CDC3-V<sub>N</sub>::TRP1*) (Oh et al., 2013).
<sup>c</sup> A standard PCR-mediated tagging of *V<sub>C</sub>::TRP1* at the C terminus of Cdc11 was performed using pFA6a-VC-TRP1 with the primers: oCDC112 (5'-GAGAAAGAAGAAATAAGTGAGGAAGCCAAAAGCGG ACTCAGAATTCGAGCTCGTTTAAAC-3') and oCDC113 (5'-AGCCAGGTTGGAAAAAGAGGCG AAAATCAAACAGGAAGAACGGATCCCCGGGTTAATTAA-3')
<sup>d</sup> A standard PCR-mediated tagging of *V<sub>C</sub>::TRP1* at the C terminus of Cdc11 was performed using pFA6a-VC-TRP1 with the primers: oCDC112 (5'-GAGAAAGAAGAAATAAGTGAGGAAGCCAAAAGCGG ACTCAGAATTCGAGCTCGTTTAAAC-3') and oCDC111 (5'-AGCATCCGATATGCACGGGCAA AGCACTGGCGAAAATAACCGGATCCCCGGGTTAATTAA-3')
<sup>e</sup> A standard PCR-mediated tagging at the C terminus of Cdc11 was performed using pFA6a-tdTomato-NAT as a template with the primers oCDC112 and oCDC113.

**Supplementary Table S2.**
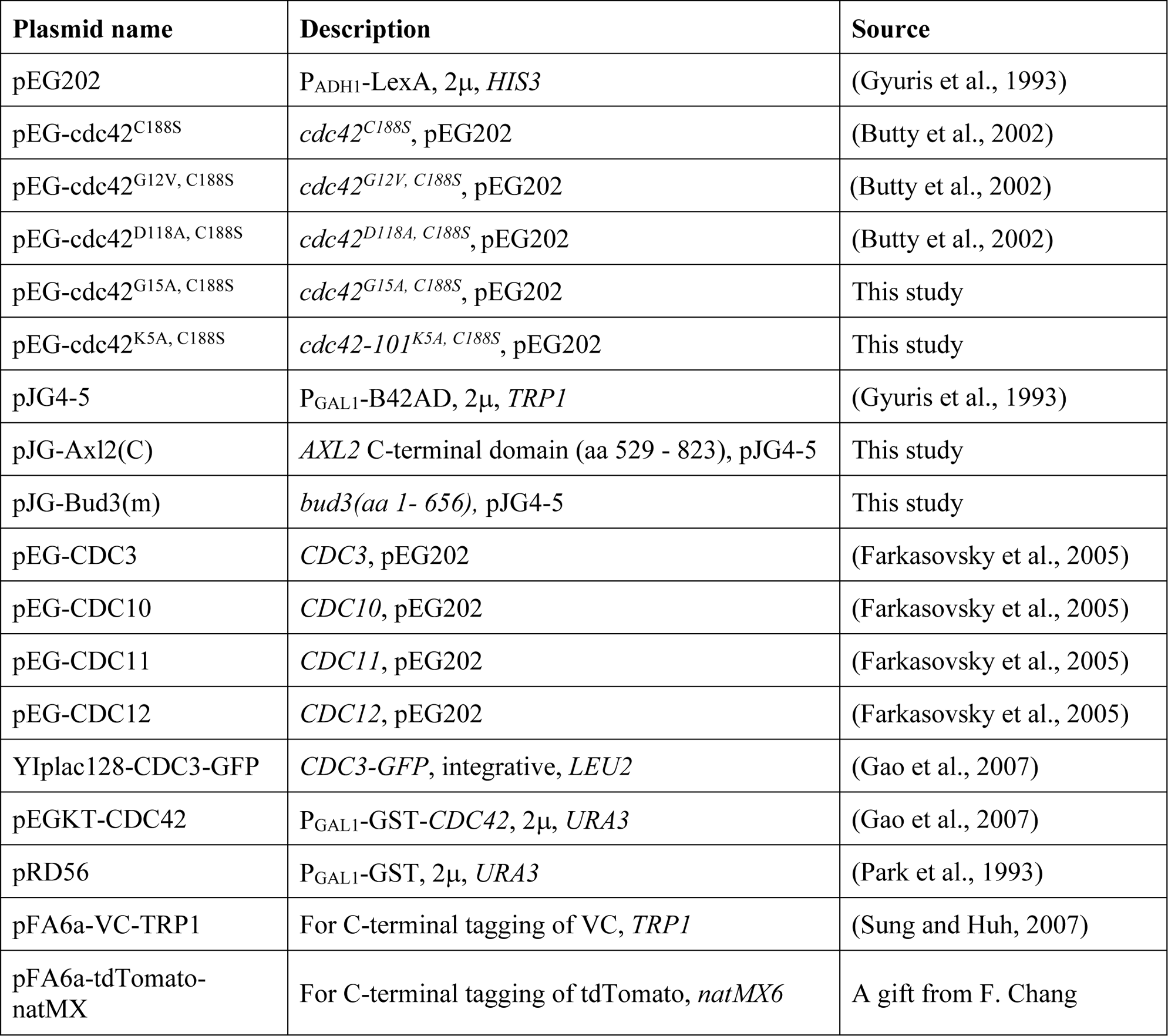
Plasmids used in this study.

## Notes

### Competing Interest Statement

The authors have declared no competing interest.

